# A conformationally heterogeneous bending pivot enables bent-to-straight transition in the central helix of mycobacterial FtsZ

**DOI:** 10.1101/2025.10.28.685025

**Authors:** Anu Sodhi, Sneh Lata, Sapana Rani, Lucky Singh, Jagrity Choudhury, Barnali Chaudhuri

## Abstract

Conformational changes in the central helix at the inter-domain cleft of bacterial treadmilling motor protein FtsZ are coupled to polymerization. Central helix of mycobacterial FtsZ interconverts between a bent and a straight form, with an unknown mechanism. We probed the mechanism of this conformational switching in the central helix of mycobacterial FtsZ using multi-temperature synchrotron crystallography at 20 ºC, 30 ºC, 37 ºC and −173 ºC temperatures. A comparison of the resultant crystal structures of FtsZ revealed altered conformations at the bending pivot of the bent central helix inside the inter-domain cleft. Further, ensemble modeling of FtsZ structure shows that this bending pivot is labile at near-physiological temperatures. Conformational fluctuations in this pivot region resulted in breakage of regular alpha helical hydrogen bonds that likely made the central helix easily bendable. These fluctuations are largely arrested in the straightened form of the central helix in comparison to the bent form. To summarize, multi-temperature crystallography combined with ensemble modeling suggest that conformational heterogeneity and associated perturbations of helix-forming interactions in the bending pivot can trigger bent-to-straight conformational transition in the central helix of mycobacterial FtsZ. This work demonstrates the effectiveness of multi-temperature crystallography in delineating the mechanisms of conformational changes in protein machines.

## Introduction

Conformational dynamics and coupled conformational changes are known enablers of protein machine function (Bustamante et al., 2001; Wei et al., 2016; Yabukarski et al., 2022). Therefore, a knowledge of protein conformational dynamics is critical for mechanistic understanding of molecular motor action, such as treadmilling, that relies on conformational changes. Filament-forming, treadmilling protein FtsZ is an integral component of the bacterial cell division machinery and is a target for antimicrobial drug discovery efforts (Du and Lutkenhaus, 2019; McQuillen and Xiao, 2020; Pradhan et al., 2021). Interconversion between a monomeric relaxed conformation, and a tensed, filament-compatible conformation, is necessary for sustaining the structural polarity of treadmilling FtsZ filament (Wagstaff et al., 2017; Wagstaff et al., 2023; Corbin and Erickson, 2020). However, detailed mechanisms of conformational changes in FtsZ are little understood.

Crystal structures of FtsZ revealed two domains, an N-terminal GTP-binding domain (NTD) and a C-terminal GTPase activating domain (CTD), that are separated by a conserved central helix housed in the inter-domain cleft (Lowe and Amos, 1998; Leung et al. 2004; Wagstaff et al., 2017; Chakraborty et al., 2024, **Figure 1A**). In addition, FtsZ contains a flexible C-terminal segment of variable length that is critical for its function (Buske and Levin, 2013). The “synergy loop” at the C-terminal end of the central helix participates in the polymerization of FtsZ, and completes the GTPase active site formed at the polymerization interface (Matsui et al., 2012; Ruiz et al., 2022; Fujita et al., 2023). A comparison of the monomeric and filamentous forms of FtsZ from *Staphylococcus aureus* (saFtsZ) showed that the central helix in the inter-domain cleft is longitudinally shifted, which is coupled with domain structure reorganization, for polymer formation (**Figure 1B**; Matsui et al., 2012; Wagstaff et al., 2017). In contrast, FtsZ from human pathogen *Mycobacterium tuberculosis* (tbFtsZ) contains a bent central helix, that adopts a more straightened conformation in a reported curved filament form (**Figure 1C**; Leung et al. 2004; Respicio et al., 2008; Li et al., 2013; Guan et al., 2018; Alnami et al., 2021). Little is known about the mechanism of this species-specific bent to straight conformational switching of the central helix observed in the crystal structures of tbFtsZ.

**Figure 1.**
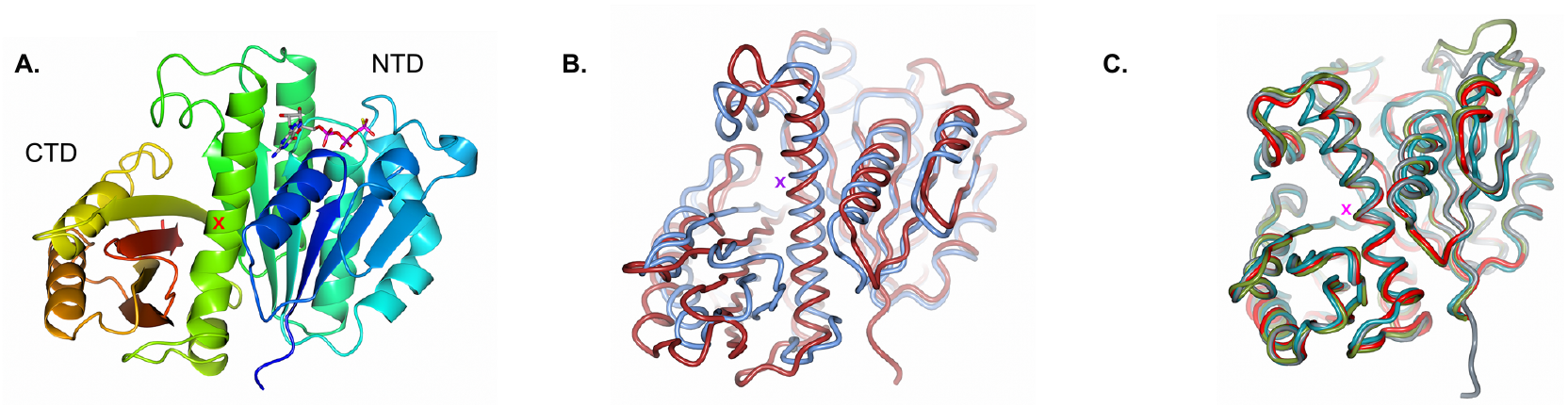
Crystal structures of FtsZ in different conformations. (A) Cartoon diagram (Jones rainbow, blue at the N-terminus and red at the C-terminus) of a crystal structure of tbFtsZ (pdb code: 1rlu). A GTP-γ-S molecule bound to the active site in the NTD is shown as a stick. (B) Ribbon diagrams of saFtsZ in two conformations (pdb codes: 5mn5 in blue and 3voa in red). (C) Ribbon diagrams of crystal structures of tbFtsZ (pdb codes: 2q1x A chain in red, 1rq2 B chain in blue, 5zue A chain in grey and 4kwe A chain in olive). The central helix region adopts either a bent (1rq2 B chain) or a straight form (2q1x A chain, 5zue A chain, 4kwe A chain). Positions of the central helices in all figures are marked with an X sign in orange.

Comparison of crystal structures of motor proteins in different conformations traditionally yielded critical insights on the mechanisms of molecular motor actions (Wagstaff et al., 2017; Wagstaff et al., 2023). However, majority of structures in PDB are determined from cryo-cooled crystals, which are dominated by protein conformations below glass transition temperature (Keedy, 2019; Fischer, 2021; Thorne, 2023). Structures determined using diffraction data collected at multiple temperatures, particularly at temperatures above glass transition temperatures, can provide a better understanding of physiologically relevant conformational fluctuations in the active site or allosteric cleft or hinge regions (Frauenfelder et al., 1979; Fraser et al., 2011; Keedy et al., 2015; Keedy, 2019; Keedy et al., 2018; Fischer, 2021; Thompson, 2023; Greisman et al., 2023; Brink et al., 2025). Shrinked unit cells due to flash-cooling and associated repacking, presence of cryoprotectants and other factors can have an effect on protein conformations, that can conceal biologically relevant conformational states (Thorne, 2023). It was known for a long time that crystallography can reveal alternative conformational states in proteins (Smith et al., 1986). Non-cryogenic crystallography at multiple temperatures can unveil those functionally relevant “hidden alternative states” that can be missed because of conformational rearrangements in cryo-cooled crystals (Fraser et al., 2009; Keedy et al, 2009). Multi-temperature crystallography as an experimental approach holds great promise for understanding the conformational changes that are indispensable for proper functioning of molecular motors, such as FtsZ.

Many would argue that multi-conformer or ensemble model of a protein structure, and not the traditional single conformer model, can better describe protein function in terms of structural dynamics (Furnham et al., 2006; Levin et al., 2007; Woldeyes et al., 2014). History of method development for modeling protein conformational dynamics in crystal structures is long and arduous (Gros et al., 1990; Kuriyan et al., 1991; Pellegrini et al., 1997; DePriesto et al., 2004; Terwilliger et al., 2007, van den Bedem et al., 2009; Woldeyes et al., 2014; Ebrahim et al., 2022; Wankowicz et al., 2024). Advances in methods for modeling low-occupancy rotamers and other developments showed promise for detecting lowly populated conformational states that are critical for protein function (Riley et al., 2021; Stachowski and Fischer, 2023; Wankowicz et al., 2024). Recently, an ensemble refinement method (ER) that use molecular dynamics simulation restrained by X-ray diffraction data showed that ensemble modeling can be possible even at a resolution as low as about 3.0 Å (Burnley et al., 2012; Forneris et al., 2021; Ploscariu et al., 2021). These ensemble models are better supported by NMR dipolar residual coupling data (Shen et al., 2023). Ensemble modeling, coupled with multi-temperature crystallography, are emerging as new ways to characterize functionally relevant conformational dynamics in protein machines.

Here, we report five crystal structures of tbFtsZ at 20 ºC (20C-tbFtsZ-I and 20C-tbFtsZ-II), at 30 ºC (30C-tbFtsZ-I), at 37 ºC (37C-tbFtsZ-I), and at −173 ºC (cold-tbFtsZ-I) that suggests a mechanism of bent-to-straight conformational switching of the central helix. These structures are the first non-cryogenic models of tbFtsZ, including a structure of tbFtsZ at human body temperature, which is the natural ambient temperature for *M. tuberculosis*. The inter-domain cleft of FtsZ is an allosteric binding site for inhibitors (Pradhan et al., 2021). A comparison of our reported tbFtsZ structures revealed a modified inter-domain cleft that partly blocks a previously known inhibitor binding site.

Analysis of these crystal structures of tbFtsZ determined at different temperatures showed functionally relevant conformational changes in the central helix. Ensemble modeling of tbFtsZ structures at 20 ºC and 30 ºC suggests that conformational fluctuations in the backbone atoms in the bending pivot region are perturbing regular alpha-helical hydrogen bonds. The pivot of helix bending showed significant heterogeneity in the bent central helix, that was mostly reduced in the straightened helical form. As a result, the central helix can undergo significant conformational changes centered at the bending pivot within the cleft that ranges from bent to straight helical forms. To summarize, multi-temperature crystallography combined with ensemble modeling provided a mechanistic understanding of conformational switching in the central helix of tbFtsZ in terms of structural heterogeneity.

## Results

### Multi-temperature crystallography shows conformational heterogeneity in the inter-domain cleft of tbFtsZ

A comparison of the single conformer models of tbFtsZ showed that major conformational changes were mainly localized in a few exposed loop regions, and in the inter-domain cleft. One of the two chains of tbFtsZ (the A chain) in the asymmetric unit contain a relatively straight central helix in the cleft, which is similar to the reported curved filamentous form of tbFtsZ (**Figure 1C**; Li et al., 2013). This straight form of central helix adopt comparable conformations in all our tbFtsZ structures across different temperatures (**Figure 2A**). On the other hand, the B chain of tbFtsZ contains a bent central helix (**Figure 1C**), which showed noticeable conformational variability amongst our tbFtsZ structures (**Figure 2B**). Furthermore, the isomorphous difference density maps between 20C-tbFtsZ-I and 30C-tbFtsZ-I, and between 20C-tbFtsZ-I and cold-tbFtsZ-I, suggest higher level of variability in the bent helix than the straight helix form of tbFtsZ (**Figure 2A-B**). Crystal structure of the cryo-protected tbFtsZ is somewhat dissimilar to the tbFtsZ structures at physiological and near-physiological temperatures (**Figure 1C**).

**Figure 2.**
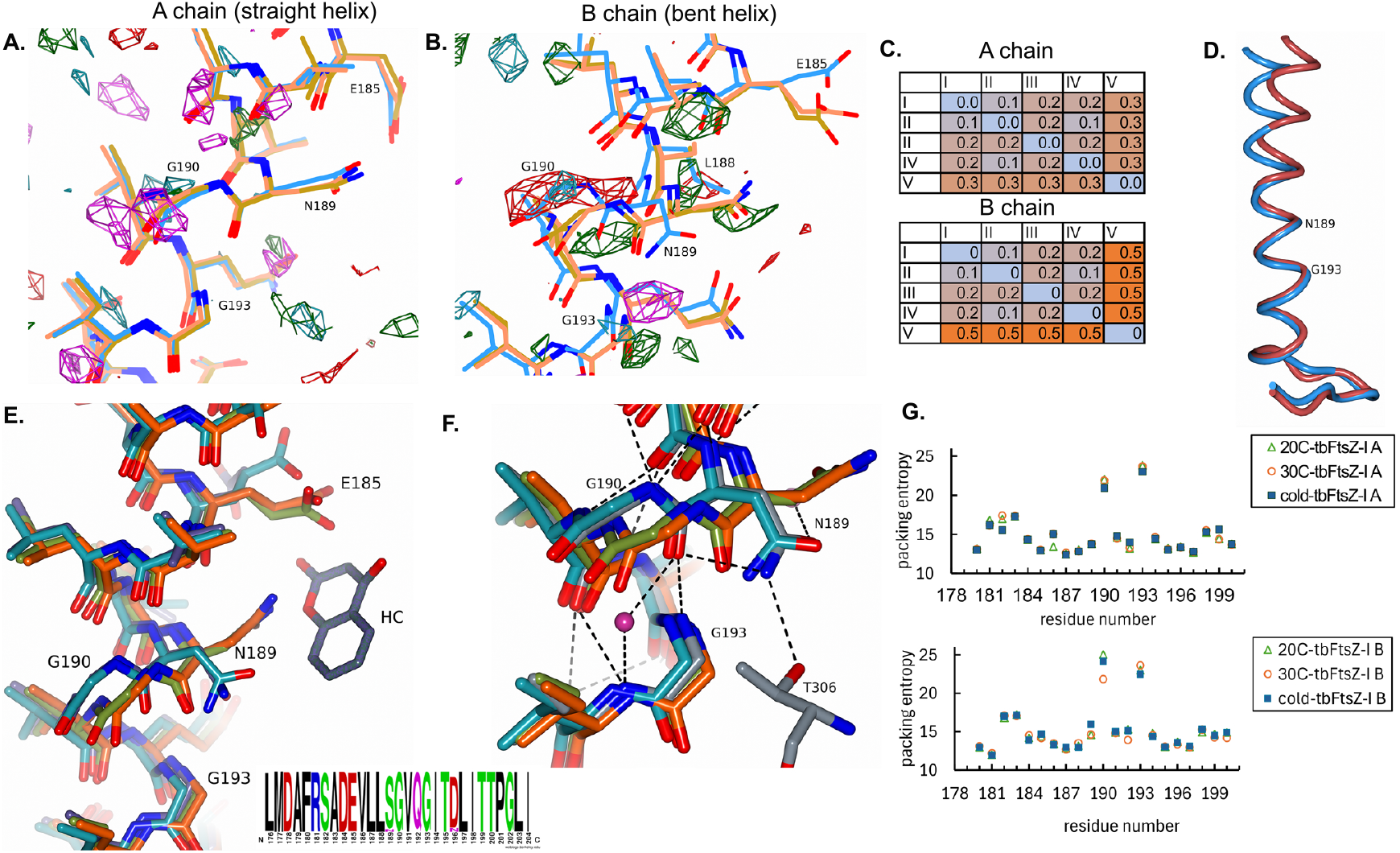
Conformational heterogeneity in the central helix of tbFtsZ. (A-B) Central helix regions of the single conformer models of tbFtsZ in the A chain (A) and B chain (B) are shown as sticks (20C-tbFtsZ-I in brown, 30C-tbFtsZ-I in crimson and cold-tbFtsZ-I in blue), along with the fo-fo maps (fo is the observed structure factor). The fo-fo maps between 20C-tbFtsZ-I and 30C-tbFtsZ-I and between 20C-tbFtsZ-I and cold-tbFtsZ-I (calculated using the coordinates of 20C-tbFtsZ-I) are contoured at +-3 sigma, and are shown with following color scheme: positive density dark green and negative density red for 20C-tbFtsZ-I and cold-tbFtsZ-I; positive density dark cyan and negative density magenta for 20C-tbFtsZ-I and 30C-tbFtsZ-I. Oxygen and nitrogen atoms are shown in red and blue color in all figures. (C) Heatmap representation of the root mean squared deviation (RMSD, in Å) between the single conformer models of 20C-tbFtsZ-I (I), 20C-tbFtsZ-II (II), 30C-tbFtsZ-I (III), 37C-tbFtsZ-I (IV) and cold-tbFtsZ-I (V) are shown for the A chains (top panel) and B chains (bottom panel). (D) Superposed central helix of the A chain (blue) and B chain (red) of 20C-tbFtsZ-I are shown as worms. Cα atoms of residues 193 to 208 were used for structural superposition. (E) Central helix segments of the B chains of tbFtsZ are shown in stick representation for 20C-tbFtsZ-I (green), 30C-tbFtsZ-I (orange), cold-tbFtsZ-I (blue), along with the hydroxycoumarin-bound tbFtsZ (pdb code 6y1u; grey). Hydroxycoumarin (HC) is shown as a stick in purple. In the inset (lower right), a sequence logo depicting the degree of sequence conservation in the central helix of actinobacterial FtsZ (residues 176-204) is shown. (E) A close up view of the residues 189 to 194 in the B chains of the single conformer models of 20C-tbFtsZ-I (green), 30C-tbFtsZ-I (orange), cold-tbFtsZ-I (blue), and a high resolution structure of tbFtsZ (pdb code 6ym9; 2.0 Å, grey). Hydrogen bonds formed in the high resolution structure of tbFtsZ (6ym9), including those with bound water molecules (red ball) and T306 residue from the CTD (grey), are shown as black, dashed lines (oxygen: red; nitrogen: blue in A, B, E and F). (F) Plot of packing entropy *versus* residue number in the central helix regions of the A chain (upper panel) and B chain (lower panel) of the single conformer models of tbFtsZ.

The pivot of helix bending that encompass 189-193 residues in the central helix of tbFtsZ contain two glycine residues (G190 and a highly conserved G193; **Figure 2D-F**). Even though not conserved across all bacterial FtsZ, G190 is highly conserved in actinobacterial FtsZ (**Figure 2E**). Packing entropy of both G190 and G193 residues are higher than the surrounding residues inside the cleft, suggesting the availability of room for conformational flexibility in this region (**Figure 2G**). In the previously reported high resolution cryo-cooled crystal structures of tbFtsZ, and in cold-tbFtsZ-I, the G190 carbonyl group of the B chain points towards a hydrophobic wall of the cleft lined by the side-chains of F97, I159, I194 residues (Alnami et al., 2021). A recent analysis of protein crystal structures determined at ultra-high resolution suggests that main chain carbonyl oxygen atoms of alpha helices can be in protonated form (Panjikar and Weiss, 2025). Such protonation state of the carbonyl moiety of G190 probably allowed it to move out towards a hydrophobic part of the cleft. A water molecule occupies the space between the backbone atoms of G190 and I194 in the bent central helix (Alnami et al., 2021; **Figure 2F**). This water molecule was not observed in the structure of cold-tbFtsZ-I, which is otherwise similar to the previously known cryo-protected structures of tbFtsZ.

In the structures of tbFtsZ that were determined at 20 ºC and 30 ºC, the G190 carbonyl group of the B chain (bent helix form) does not participate in regular “helix-forming” hydrogen bonding interaction (**Figure 2E-F**). The Cα atom of this G190 residue is shifted by more than 1.5 Å, with a concomitant change in the backbone conformation (Δϕ ∼ 23.5 º, ΔΨ ∼ 2 º), between 20C-tbFtsZ-I and cold-tbFtsZ-I structures (**Figure 2B, E-F; Figure S1**). The pivot of the bent helix in cryo-protected cold-tbFtsZ-I is likely in an annealed conformation (**Figure 2E-F**). The backbone atoms of tbFtsZ at 20 ºC and 30 ºC adopt distinct conformations in the pivot region of the bent central helix when compared to the backbone of cold-tbFtsZ-I.

Varied rotameric conformations were observed for E185, L188 and N189 residues in the bent helix of tbFtsZ (**Figure 2A-B, E-F, Figure S1C**). Although the E185 and L188 residues are conserved in actinobacteria, the N189 residue is often replaced by a serine residue with a shorter side-chain (**Figure 2E**). Hydrogen bonding interaction between the side-chains of N189 and T306 from the CTD across the cleft was not formed due to repositioning of N189 side chain in 20C-tbFtsZ-I and 30C-tbFtsZ-I (**Figure 2F**). Altered rotamers of N189 and E185 partly occlude a previously reported hydroxycoumarin binding site within the cleft of tbFtsZ (**Figure 2E**). As a result of these conformational changes, accessibility of the cleft in the B chain of tbFtsZ was noticeably modified.

### Ensemble modeling of tbFtsZ revealed backbone conformational fluctuation within the bending pivot of the central helix

To further probe heterogeneity in the tbFtsZ structures, we used recently developed Ensemble Refinement (ER) approach (Burnley et al., 2012; **Figure 3A-D; Figure S2**). As anticipated, resultant structures in the ensembles of tbFtsZ are quite heterogeneous in the region surrounding the N189 and G190 residues in the bending pivot (**Figure 3A-D, 4A-D**). Backbone and side-chain atoms of N189 showed high positional variability, which was particularly pronounced in the B chain (**Figure 4; Figure S2**). The backbone nitrogen of G193 forms a hydrogen bond with the backbone carbonyl group of N189. The G190 residue, and not the conserved G193 residue, showed drastic conformational variation in the ensemble structures (**Figure 4**). Kinetic energy of the pivot residues were some-what higher than that of the surrounding residues in the ensemble models of the B chain, which is probably due to high mobility (**Figure 3**).

**Figure 3.**
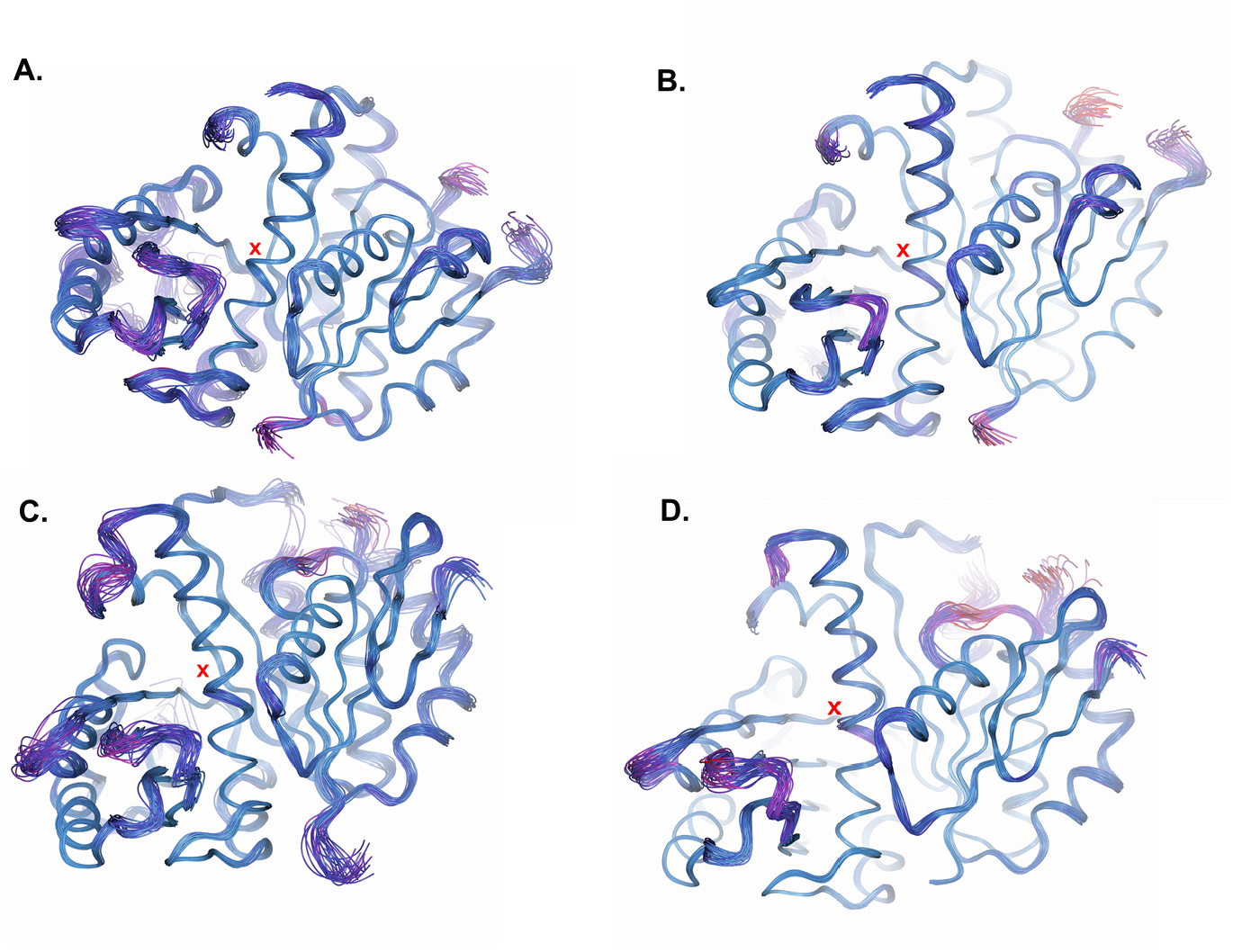
Ensemble models of tbFtsZ at 20 ºC and 30 ºC. (A-D) Ensemble models of 20C-tbFtsZ-I A chain (A), 30C-tbFtsZ-I A chain (B), 20C-tbFtsZ-I B chain (C) and 30C-tbFtsZ-I B chain (D), colored by kinetic energy (low to high: blue to magenta), are shown as worms. Positions of the bending pivots (near 189^th^ residue) are marked with red X signs in A-D.

**Figure 4.**
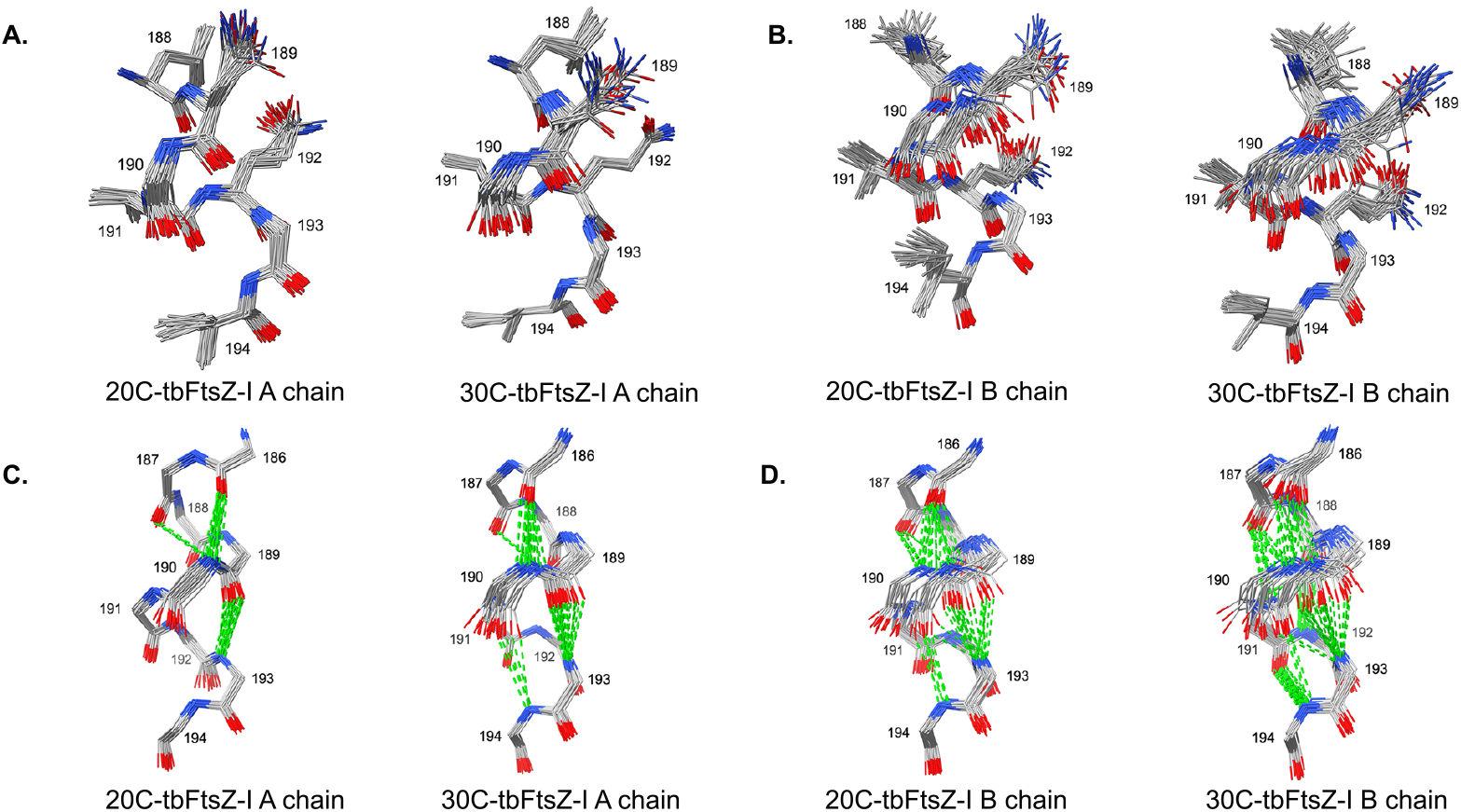
Ensemble modeling shows conformational variations at the bending pivot. (A-B) Residues 188-194 of the A chains (A) and B chains (B) of the ensemble models of 20C-tbFtsZ-I (left)) and 30C-tbFtsZ-I (right) are shown in stick representation. (C-D) Backbone atoms of residues 186-194 of the A chains (C) and B chains (D) of 20C-tbFtsZ-I (left) and 30C-tbFtsZ-I (right) are shown as sticks along with hydrogen bonds (green dashed lines) formed by glycine residues (G190 and G193). Side chains are not shown in C and D for clarity. (carbon in grey; oxygen in red; nitrogen in blue)

Conformational heterogeneity in the backbone atoms of the pivot residues affected their capacity to form regular alpha helical i to i+4 hydrogen bonds (**Figure 4**). The main-chain polar atoms of the pivot residues in these ensembles can be classified into two states. Some of these are in “helix-forming” state, with a regular alpha-helical hydrogen bond between the backbone carbonyl groups and nitrogen atoms (**Figure 4A-B**). Others are in a “helix breaking” state, in which the canonical i to i+4 hydrogen bonds are not formed (**Figure 4A-B**). Ensemble modeling suggests that movement of the central helix is likely caused by a labile bending pivot, in which the polar backbone atoms can easily interconvert between the “helix-forming” and “helix-breaking” states at ambient temperature.

### Crystal structure of tbFtsZ at the mammalian body temperature

Next, we determined the crystal structure of tbFtsZ at 37 ºC (37C-tbFtsZ-I), which is the physiologically relevant temperature for proteins from human pathogen *M. tuberculosis*. Due to modest resolution, detailed conformational dynamics of 37C-tbFtsZ-I could not be modeled using ensemble refinement approach. Final refined single conformer model of 37C-tbFtsZ-I was compared with another single conformer model of tbFtsZ at 20 ºC (20C-tbFtsZ-II), both of which were determined at an equivalent resolution at different temperatures (**Figure 5A-B**). Structural superposition shows that the structural models of 37C-tbFtsZ-I and 20C-tbFtsZ-II are largely similar (∼ 1 Å for 585 Cα atoms from both chains). Since we did not obtain radiation damage-free temperature-factors, the temperature-factors were not analyzed. No additional level of disorder, such as a molten core (Burnley et al., 2012), was observed in the difference electron density map associated with the structure of 37C-tbFtsZ-I (**Figure 5A-B**). Similarity between the structures of tbFtsZ at human body temperature and at room temperature suggest that a room temperature model can be useful for learning about the functional conformation of tbFtsZ at human body temperature.

**Figure 5.**
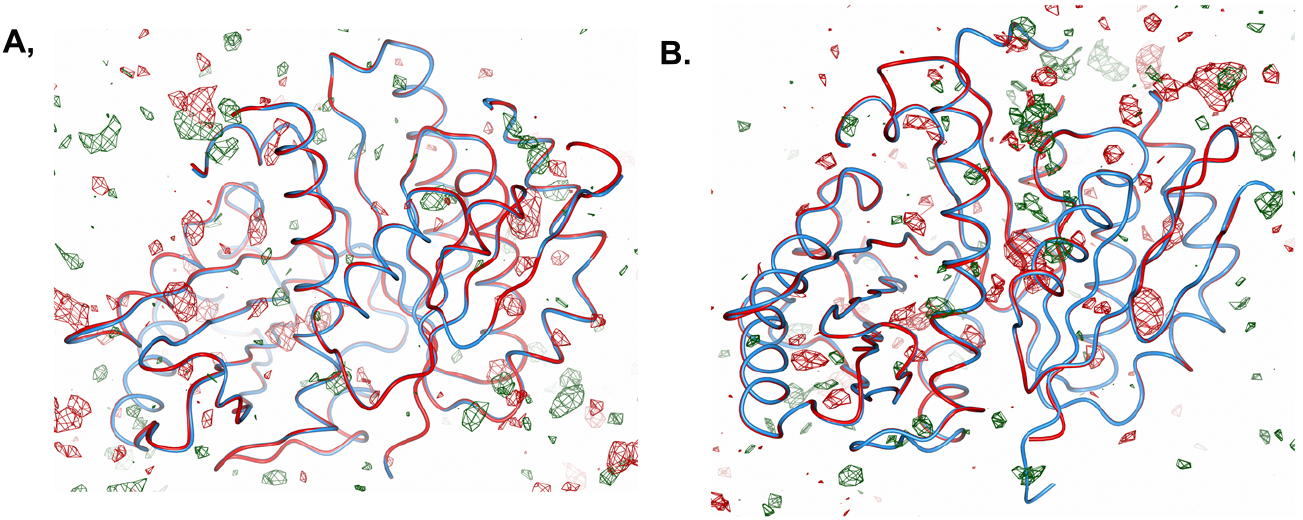
Crystal structures of tbFtsZ at physiologically relevant temperatures. (A-B) Cα traces of 37C-tbFtsZ-I (red worm) and 20C-tbFtsZ-II (blue worm), along with the mfo-Dfc map associated with 37C-tbFtsZ-I model, are shown for the A chain (A) and B chain (B). The mfo-Dfc maps (contoured at ± 3 σ) are shown in A-B with following color scheme: positive density in dark green, negative density in red.

## Discussion

Multi-temperature crystallography of tbFtsZ elegantly revealed that the pivot of the bent helix is highly labile when compared to the pivot of the straight helix (**Figure 2, 4, 6**). What purpose does this labile pivot located in the middle of a sheltered cleft serve? It appears that the pivot regulates the movement of the central helix. Regulation of movement is achieved by fluctuation-driven disruptions of helix-stabilizing interactions in the pivot (**Figure 4**). A large number of breakage or distortion of hydrogen bonds in the bending pivot effectively permits the central helix to sample wider conformational space, including the more labile bent forms and the relatively rigidified straightened forms (**Figure 4, 6**).

**Figure 6.**
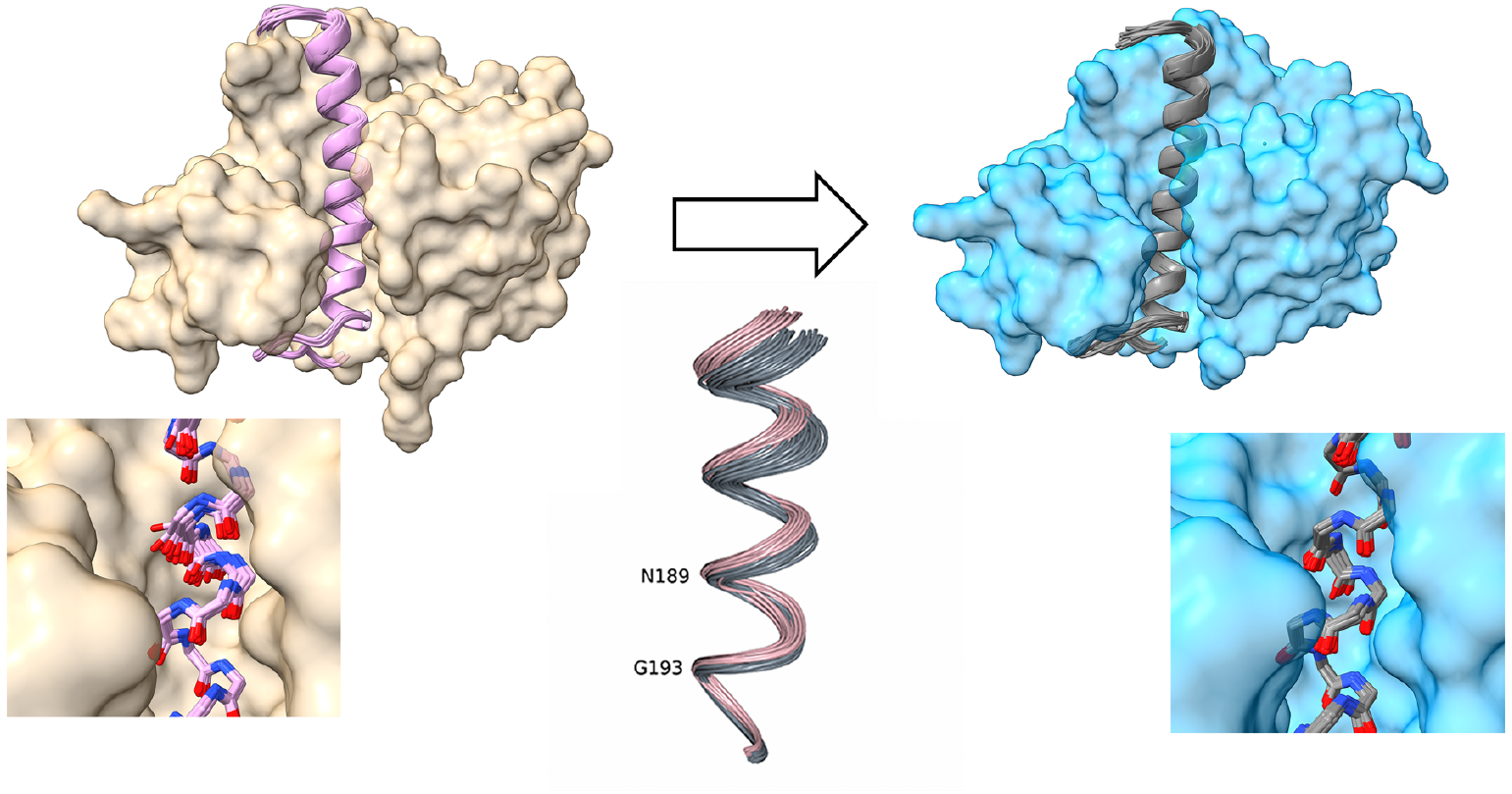
Heterogeneity in the bending pivot enables bent-to-straight conformational transition in the central helix. Upper panel, left: surface of the B chain of tbFtsZ (pale yellow, single conformer) excluding the central helix, along with the bent central helix segments of the ensemble model (pink cartoon). Upper panel, right: surface of the A chain of tbFtsZ (blue, single conformer) excluding the central helix, along with the straight central helix segments of the ensemble model (grey cartoon). Lower panel: backbone atoms of the bending pivot region are shown as sticks, along with the surfaces, in the same way as shown in the upper panel. In the center, central helices of the ensemble models (A chains in grey, B chains in pink) are shown as worms. Single conformer and ensemble models of 20C-tbFtsZ-I are shown in all panels.

Even at modest resolutions, the presented structures of tbFtsZ clearly showed conformational variations in the bending pivot (**Figure 2, Figure S1**). Structures free of radiation damage that can be potentially obtained from “diffraction before destruction” mode of data collection (Neutze et al., 2000, Chapman et al., 2014) were not acquired due to a lack of access to such data collection facilities. However, according to a previous study, conformational variations observed in ambient temperature crystal structures are shown to be only mildly influenced by radiation damage (Russi et al., 2017). Therefore, these observed conformations that are supported by electron density maps appear to be genuine features of tbFtsZ structures (**Figure S1**). Since modeling alternative conformations require higher resolution, we used newly developed ER method (Burnley et al., 2012) to model structural heterogeneity. This study shows that multi-temperature crystallography conducted using a second generation synchrotron X-ray source can yield useful insights about functionally relevant conformational fluctuations in a molecular machine.

In the crystal structures of tbFtsZ at 20 ºC and 30 ºC, the inter-domain cleft housing the bent helix have an altered accessibility to inhibitors. Changed rotamer conformations in the cleft partially occlude a hydroxycoumarin binding site that was found in a previously reported cryo-protected structure of tbFtsZ (**Figure 2E;** Alnami et al., 2021). This finding is in line with a previous large scale study in which comparison of ligand binding showed that binding mode can be changed in the room temperature structures in comparison to the cryo-cooled structures (Mehlman et al., 2023). In saFtsZ, the inter-domain cleft is the binding site of an allosteric inhibitor PC190723 (Matsui et al., 2012). This cleft in FtsZ is a target for numerous structural studies, and structure-guided drug discovery efforts (Pradhan et al, 2021; Poddar et al., 2025). Our results suggest that crystal structures of FtsZ determined at multiple, physiological and near-physiological temperatures can be more suitable starting points for structure-based allosteric inhibitor design efforts with an aim to dampen conformational fluctuations at the cleft.

Even though a crystal structure of curved filamentous form of tbFtsZ has been reported (Li et al., 2013), structure of a straight, filamentous form of tbFtsZ is not available. A low resolution (∼ 7 Å) cryo-EM map of tbFtsZ filament appears to fit a filamentous form of saFtsZ (Wagstaff et al., 2023). It is possible that the central helix in tbFtsZ will undergo further conformational change for filament formation, which will be likely complemented by global change in domain organization. Dynamics of the central helix can play a critical role in conformational shifting and longitudinal interface formation required for tbFtsZ polymerization.

Nucleotides are essential ingredients for treadmilling in proteins (Narita, 2011). How nucleotide binding affects conformational switching of FtsZ is much discussed in the literature (Corbin and Erickson, 2020; Wagstaff et al., 2023). Binding of a small molecule, such as a nucleotide, can potentially redistribute conformations within an ensemble to facilitate protein action (Wancowicz and Fraser, 2025). The D184 residue from the N-terminal end of the central helix in tbFtsZ interact with the bound nucleotide in the active site (Leung et al. 2004). Therefore, movement of the central helix will certainly get affected by a bound nucleotide, and *vice versa*. In spite of numerous co-crystallization and soaking attempts, we were unable to obtain any structure of tbFtsZ in complex with a nucleotide. It will be interesting to find out how nucleotide-binding in the active site of tbFtsZ, and hydrolysis, affects conformational dynamics of the central helix and how this modulation connects to tbFtsZ polymerization.

## Materials and Methods

### Diffraction data collection at multiple temperatures

Expression, purification and crystallization procedures of tbFtsZ have been described elsewhere (Leung et al., 2004; Respicio et al., 2008; Alnami et al., 2021; Singh, 2023; Choudhury and Chaudhuri, 2025). Multi-temperature diffraction datasets were collected at beamline PX-BL21, Indus-2 synchrotron, Raja Ramanna Center for Advanced Technology, Indore (Kumar et al., 2016). MicroRT capillaries (Mitegen, cat number RT-T1) were used for non-cryogenic data collection. Due to the problem of severe radiation damage, a large number of crystals were tested. Ascorbic acid and beta-mercaptoethanol were used for mitigating radiation damage. Finally, diffraction data at three temperatures (20 ºC, 30 ºC and 37 ºC) were collected (**Table S1**). Data processing was performed using XDS and MOSFLM from the CCP4 suite of programs (Kabsch et al., 2010; Agirre et al., 2023).

A diffraction dataset at −173 ºC was collected at beamline ID23-2, European Synchrotron Radiation Facility (ESRF), Grenoble, France, and processed on-site (**Table S1**).

### Refinement of single conformer models of tbFtsZ

Following rigid body minimization or Phaser run in Phenix (Liebschner et al., 2019) using a known structure of tbFtsZ (PDB code 6ym9), or the final refined model of 20C-tbFtsZ-I, PDB_REDO server (Joosten et al., 2014) was used for automated rebuilding and refinement of the models. Final resolutions for refinement were decided based on paired-refinement option of PDB_REDO (Malý et al., 2020). Next, the models were manually inspected and rebuilt in Coot (Emsley and Cowtan, 2004) and further refined in Phenix using positional minimization and simulated annealing slow cool protocols. Since the two chains of tbFtsZ in the asymmetric unit adopt different conformations, non-crystallographic symmetry restraints were not applied. Secondary structure restraints were applied. Individual isotropic B-factors were refined for all structures refined at better than 3.0 Å resolution. For the two models refined at 3.2 Å resolution, grouped B-factors were refined. A citrate from crystallization buffer, which was bound to the A chain of tbFtsZ, was modeled based on the model of a similarly bound citrate in a tbFtsZ structure (pdb code 2q1x). Geometry of the citrate buffer was rather poor in our medium resolution structures of tbFtsZ, and kept as close as possible to the model from the previously reported high resolution structure of tbFtsZ. Owing to modest resolution, alternative conformations were not modeled in these tbFtsZ structures. Refinement statistics for five structures of tbFtsZ are summarized in **Table S2**.

### Ensemble refinement of tbFtsZ

Ensemble refinements (ER; Burnley et al., 2012) were performed in Phenix for the structural models at 20 ºC and 30 ºC (**Table S3**). A number of pTLS values (0.4, 0.6 and 0.8) were tested before selecting the best one with lowest R-free. Harmonic restraints were applied to non-protein moieties. Results of those ER runs that produced higher R-free than the single conformer models are not reported. Final ER runs were repeated twice for each case.

### Structure analysis

ChimeraX (rbvi.ucsf.edu/chimerax), Jalview (https://www.jalview.org/) and CCP4 Molecular graphics (Agirre et al., 2023) were used for structure/sequence analysis and figure preparations. MUSTANG (Konagurthu et al., 2006) and PRO-FIT (http://www.bioinf.org.uk/software/profit/) were used for structure superposition. The Basic Local Alignment Search Tool (BLAST; http://blast.ncbi.nlm.nih.gov) was used for obtaining sequence homologs of tbFtsZ. 498 actinobacterial FtsZ sequences (excluding mycobacteria) were used for preparing the sequence logo encompassing the central helix using the WebLogo server (weblogo.berkeley.edu). Polder maps, composite omit maps and isomorphous difference maps were calculated in Phenix (Liebschner et al., 2019). Main chain torsion angles were calculated in DSSP (Kabsch and Sander, 1983). Packing entropy values were calculated using PACMAN webserver (Khade and Jernigan, 2023). Hydrogen bonds were calculated in ChimeraX using 0.4 Å tolerance.

## Data availability statement

Crystal structures and associated data are submitted to PDB with following accession numbers: 9XA6, 9XE3, 9XDH, 9XE2, 9XF0

Ensemble models associated with 9XA6 and 9XDH datasets are uploaded in Zenodo (10.5281/zenodo.17469883).

## Acknowledgments

BC wish to thank CSIR Fundamental and Innovative Research of tomorrow (CSIR-FIRST; MLP061) program for funding this project, and CSIR-IMTECH for providing access to the in-house facilities. We thank staffs at beamline PX-BL21 at Indus-2 synchrotron, RRCAT, Indore, for assistance during non-cryogenic data collection. Cryogenic data was collected at beamline ID23, at ESRF, France, which was supported by Department of Biotechnology (DBT), Govt. of India and Regional Centre of Biotechnology, Faridabad. AS and LS thanks CSIR for fellowship. SL thanks Indian Council of Medical Research for her fellowship. JC thanks DBT for her fellowship.

## Notes

### Competing Interest Statement

The authors have declared no competing interest.

