## Supplemental Figures S1, S2 and Tables S1, S2 and S3 for "A conformationally heterogeneous bending pivot enables bent-to-straight transition in the central helix of mycobacterial FtsZ"

**Figure S1.**

**A.** **B.**

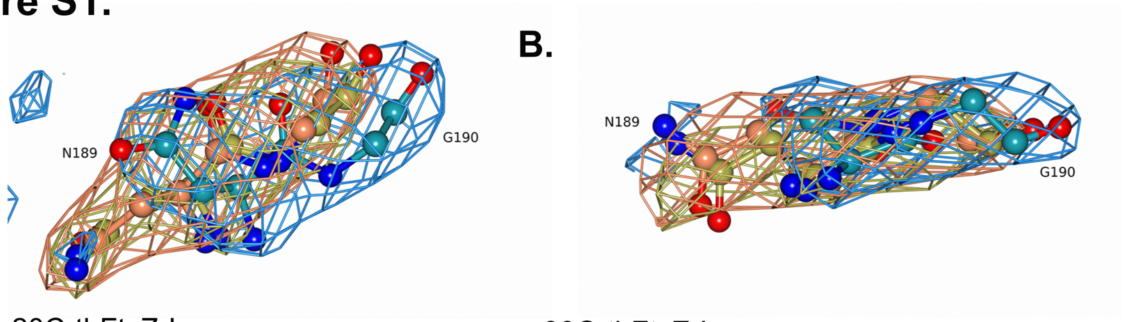


**C.** 20C-tbFtsZ-I 30C-tbFtsZ-I

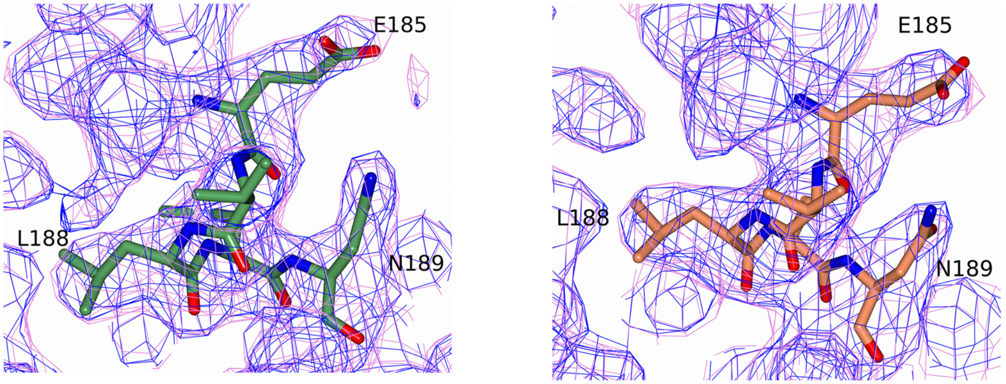


cold-tbFtsZ-I

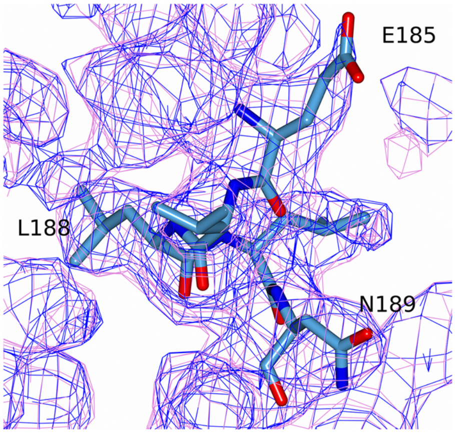


**D.**

| RSCC | E185 | L188 | N189 | G190 |
|---|---|---|---|---|
| **20C-tbFtsZ-I** | 0.94 (0.92) | 0.97 (0.94) | 0.91 (0.86) | 0.95 (0.91) |
| **30C-tbFtsZ-I** | 0.95 (0.94) | 0.97 (0.94) | 0.95 (0.89) | 0.98 (0.935) |
| **cold-tbFtsZ-I** | 0.91 (0.87) | 0.97 (0.93) | 0.96 (0.93) | 0.98 (0.95) |

**Figure S1**. (A-B) Polder omit maps (chicken-wire, contoured at +6 σ) for the N189 and G190 residues (ball-and-stick) in the B chains of 20C-tbFtsZ-I (gold for carbon and the map), 30C-tbFtsZ-I (crimson for carbon and the map) and cold-tbFtsZ-I (blue for carbon and the map) are shown. (C) Composite omit maps (orchid) and 2mfo-Dfc maps (blue) are shown (chicken-wire, contoured at 1 σ) along with the residues 185-189 (stick) from the B chains of the single conformer models of 20C-tbFtsZ-I (green), 30C-tbFtsZ-I (crimson) and cold-tbFtsZ-I (blue). Oxygen and nitrogen atoms are colored in red and blue, respectively, in A-C. (D) Real space correlation coefficients (RSCC) for E185, L188, N189 and G190 residues in the single conformer models of tbFtsZ (B chain) are shown. RSCC values were calculated in Phenix using the 2mfo-Dfc maps. RSCC calculated using the composite omit maps are in parenthesis.

### Figure S2.

**A.**

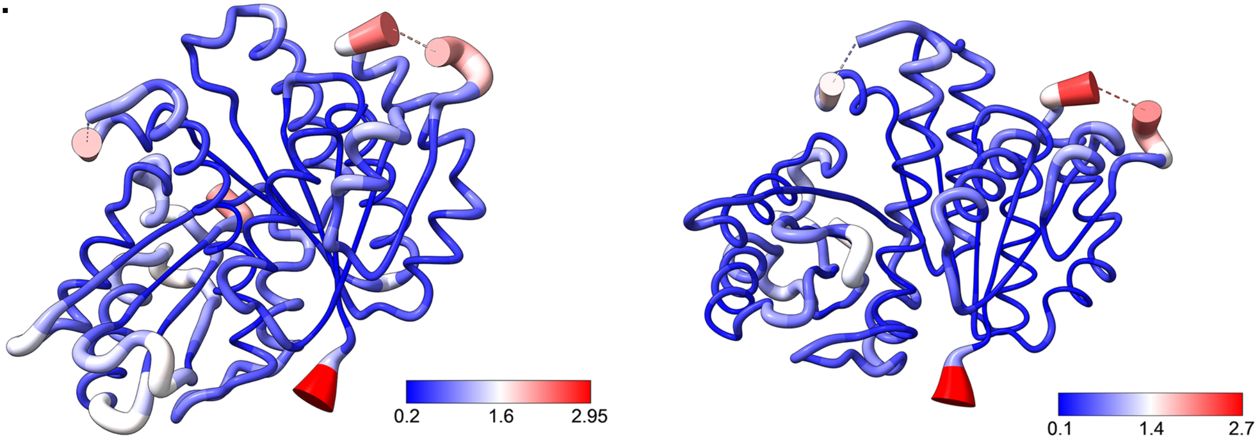


20C-tbFtsZ-I A chain

30C-tbFtsZ-I A chain

**B.**

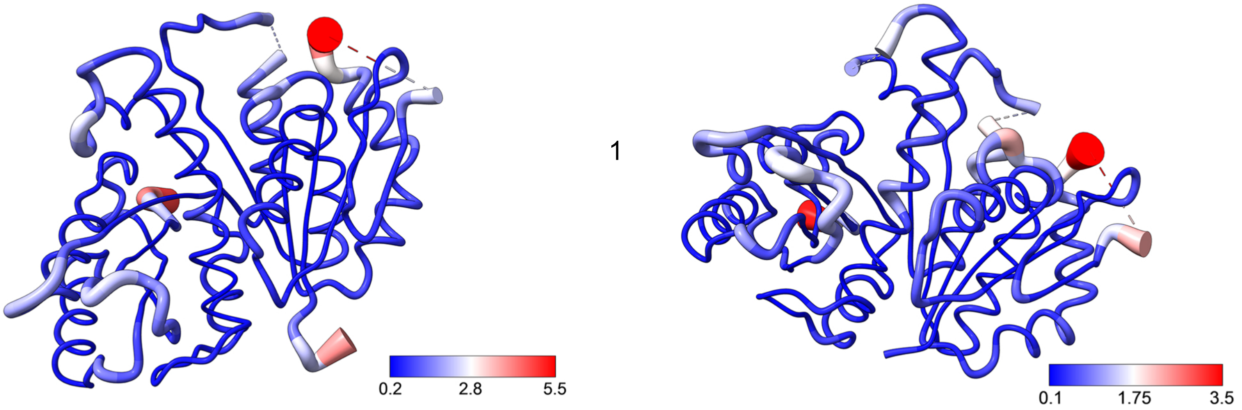


20C-tbFtsZ-I B chain

30C-tbFtsZ-I B chain

**Figure S2**. (A-B) Representative ensemble models of the A chains (A) and B chains (B) of 20C-tbFtsZ-I and 30C-tbFtsZ-I are shown as worms in putty representation, colored and scaled by RMSD based on Cα atoms over all models in the ensemble. Color bars are shown for RMSD in Å.

**Table S1**. Data collection statistics (calculated by AIMLESS, CCP4 suite of software).

| | 20C-tbFtsZ-I | 20C-tbFtsZ-II | 30C-tbFtsZ-I | 37C-tbftsZ-I | Cold-tbFtsZ-I |
|---|---|---|---|---|---|
| **Data collection temperature (°C)** | 20 | 20 | 30 | 37 | -173 |
| **Wavelength (Å)** | 0.98 | 0.98 | 0.98 | 0.98 | 0.87 |
| **Resolution range (Å)** | 78.1-2.6 (2.7-2.6) | 43.7-3.2 (3.4-3.2) | 39.6-2.8 (2.95-2.8) | 39.4 – 3.2 (3.4-3.2) | 47.9 – 2.6 (2.7-2.6) |
| **Space group** | $P6_5$ | $P6_5$ | $P6_5$ | $P6_5$ | $P6_5$ |
| **Unit cell (Å and °)** | 90.1, 90.1, 182.7, 90, 90, 120 | 90.1, 90.1, 182.8, 90, 90, 120 | 90.5, 90.5, 183.75, 90, 90, 120 | 90.3, 90.3, 182.7, 90, 90, 120 | 90.2, 90.2, 181.7, 90, 90, 120 |
| **Total reflections** | 198539 (24138) | 52761 (9597) | 79334 (11637) | 72102 (13222) | 188310 (23895) |
| **Unique reflections** | 25852 (3152) | 12769 (2337) | 20689 (3014) | 13914 (2515) | 25339 (3142) |
| **Multiplicity** | 7.7 (7.7) | 4.1 (4.1) | 3.8 (3.9) | 5.2 (5.3) | 7.4 (7.6) |
| **Completeness (%)** | 100.0 (100.0) | 92.1 (93.4) | 98.9 (99.6) | 99.7 (99.9) | 98.3 (100.0) |
| **Mean I/sd(I)** | 8.7 (2.1) | 5.1 (0.8) | 6.6 (0.6) | 4.3 (0.9) | 8.5 (1.2) |
| **Wilson B-factor** | 36.1 | 85.9 | 76.5 | 89.15 | 51.9 |
| **R-merge** | 0.33 (3.24) | 0.23 (1.59) | 0.10 (1.32) | 0.43 (2.17) | 0.19 (1.64) |
| **R-meas** | 0.37 (3.71) | 0.30 (2.075) | 0.14 (1.79) | 0.53 (2.69) | 0.22 (1.91) |

| | | | | | |
|---|---|---|---|---|---|
| **R-pim** | 0.19 (1.92) | 0.19 (1.32) | 0.09 (1.20) | 0.29 (1.56) | 0.08 (0.70) |
| **$CC_{1/2}$** | 0.98 (0.30) | 0.98 (0.29) | 0.98 (0.30) | 0.96 (0.315) | 0.99 (0.42) |

Statistics for the highest-resolution shell are shown in parentheses.

**Table S2.** Refinement statistics for the single conformer models of tbFtsZ (calculated by Phenix).

| | 20C-tbFtsZ-I | 20C-tbFtsZ-II | 30C-tbFtsZ-I | 37C-tbftsZ-I | Cold-tbFtsZ-I |
|---|---|---|---|---|---|
| **Resolution range (Å)** | 45.0 - 2.6 (2.7 – 2.6) | 39.4 – 3.2 (3.45 – 3.2) | 36.4 – 2.8 (2.9 – 2.8) | 33.1 – 3.2 (3.45 – 3.2) | 47.9 – 2.6 (2.7 – 2.6) |
| **Reflections used in refinement** | 25763 (2842) | 12713 (2545) | 20667 (2605) | 13809 (2758) | 25279 (2843) |
| **Reflections used for R-free** | 1315 (115) | 706 (143) | 1092 (142) | 750 (148) | 1291 (117) |
| **R-work** | 0.178 (0.270) | 0.194 (0.331) | 0.162 (0.323) | 0.186 (0.3015) | 0.193 (0.306) |
| **R-free** | 0.214 (0.320) | 0.244 (0.380) | 0.205 (0.369) | 0.227 (0.316) | 0.237 (0.366) |
| **Number of non-hydrogen atoms** | 4165 | 4109 | 4136 | 4040 | 4269 |
| **macromolecules** | 4141 | 4096 | 4123 | 4027 | 4189 |
| **Buffer** | 13 | 13 | 13 | 13 | 13 |
| **Solvent** | 11 | 0 | 0 | 0 | 67 |
| **Protein residues** | 595 | 589 | 589 | 587 | 598 |
| **RMS (bonds), Å** | 0.002 | 0.003 | 0.004 | 0.003 | 0.005 |
| **RMS (angles), °** | 0.7 | 0.6 | 0.55 | 0.7 | 0.7 |
| **Ramachandran favored (%)** | 98.80 | 99.48 | 99.13 | 99.30 | 99.32 |

| | | | | | |
|---|---|---|---|---|---|
| **Ramachandran allowed (%)** | 1.20 | 0.52 | 0.87 | 0.70 | 0.68 |
| **Ramachandran outliers (%)** | 0.00 | 0.00 | 0.00 | 0.00 | 0.00 |
| **Rotamer outliers (%)** | 0.00 | 0.00 | 0.00 | 0.00 | 0.00 |
| **Clashscore** | 2.42 | 2.82 | 2.18 | 3.00 | 1.92 |
| **Average B-factor (Å$^2$)** | 41.25 | 76.9 | 74.1 | 78.8 | 54.6 |
| **macromolecules** | 41.3 | 76.9 | 74.1 | 78.7 | 54.6 |
| **buffer** | 43.45 | 90.3 | 80.3 | 99.5 | 62.1 |
| **solvent** | 35.4 | - | - | - | 48.6 |

Statistics for the highest-resolution shell are shown in parentheses.

**Table S2**. Summary of ensemble refinement.

| **Dataset/model** | **20C-tbFtsZ-I** | **30C-tbFtsZ-I** |
|---|---|---|
| **No. of models** | 25 | 29 |
| **pTLS** | 0.4 | 0.8 |
| **R-work** | 0.169 | 0.159 |
| **R-free (ensemble model)** | 0.208 | 0.1935 |
| **R-free (single conformer)** | 0.214 | 0.205 |
